# Motion Sensing Superpixels (MOSES): A systematic framework to quantify and discover cellular motion phenotypes

**DOI:** 10.1101/248104

**Authors:** Felix Y. Zhou, Carlos Ruiz-Puig, Richard P. Owen, Michael J. White, Jens Rittscher, Xin Lu

## Abstract

Cellular motion is fundamental in tissue development and homeostasis. There is strong interest in identifying factors that affect the interactions of cells in disease but analytical tools for robust and sensitive quantification in varying experimental conditions for large extended timelapse acquisitions is limited. We present Motion Sensing Superpixels (MOSES), a method to systematically capture diverse features of cellular dynamics. We quantify dynamic interactions between epithelial cell sheets using cell lines of the squamous and columnar epithelia in human normal esophagus, Barrett’s esophagus and esophageal adenocarcinoma and find unique boundary formation between squamous and columnar cells. MOSES also measured subtle changes in the boundary formation caused by external stimuli. The same conclusions of the 190 videos were arrived at unbiasedly with little prior knowledge using a visual motion map generated from unique MOSES motion ‘signatures’. MOSES is a versatile framework to measure, characterise and phenotype cellular interactions for high-content screens.

## Introduction

In the development of multicellular organisms, different cell types expand and migrate to form defined organ structures. Tissue development and homeostasis require coordinated collective cellular motion. For example, in conditions such as wound healing, immune and epithelial cells need to proliferate and migrate (1–3). Deregulation of key signalling pathways in pathological conditions, including cancer, causes alterations in cellular motion properties that are critical for disease development and progression, for example leading to invasion and metastasis. Therefore there is strong biological and clinical need to study precise cellular motion characteristics in a tissue-relevant context. An example of a problem that would benefit from high-throughput computational analysis is the formation of stable boundaries between homo- and heterotypic cell populations. When two cell populations meet *in-vivo*, they often form a sharp, stable interface termed a ‘boundary’, with limited intermingling (4). In adult humans, sharp boundaries separate different types of epithelia: for example between the squamous and columnar epithelia in the esophagus, cervix and anus. Disruption of these boundaries can lead to disease. Disruption of the squamo-columnar epithelial boundary in Barrett’s Esophagus (BE) confers a 30-50 fold increased risk of esophageal adenocarcinoma (EAC) (5). Understanding how tissue dynamics relates to pathological phenotypes and how it can be affected by intrinsic and extrinsic factors is therefore a key issue.

Thus, there is a strong interest in live cell imaging for high throughput screening and analysis to identify targets and drugs that can cause or prevent abnormal cellular motion (6–8). Such applications require the extraction of quantitative biologically relevant information from complex time-lapse imaging data in a manner that allows quantification of phenotypic variation under different experimental conditions. However, such tools are currently limited. Existing experimental approaches mainly emphasise individual cell/cell interactions and single cell migration. Studies of cell population interactions and migration have been limited. Most assays use a single cell type (e.g. in a scratch wound healing assay (9)) or may not be suitable for handling high variability among replicates (e.g. due to variable cell transfection, seeding and growth rates (10, 11)) and are thus not suited to systematically analyse interactions between populations of different cell types particularly for high content screens.

A suitable computational method for studying cell population dynamics, including in a medium- or high-throughput manner, should be: (i) robust, i.e. able to handle inevitable variations in image acquisition and experimental protocol; (ii) sensitive, i.e. able to detect motion differences resulting from small changes in environment or stimuli with a minimum number of replicates; (iii) automatic, not requiring manual intervention except from the initial setting of parameters; and (iv) unbiased, able to characterise motion (e.g. as a motion ‘signature’) with minimal assumptions about motion behaviour. Existing approaches including vertex models (12, 13), differential equations (14, 15), cellular automata (16–18) and cell tracking (19–24) have successfully enabled the assessment of specific biological phenomena (such as individual cell motility or stresses between cell-cell contacts) but do not fully meet these four criteria, are limited in scope of application and are difficult to generalise to medium/high content screening. Specifically, vertex models, differential equations and cellular automata are founded on prior modelling assumptions that may not be satisfied for arbitrary timelapse acquisitions. Cell tracking methods rely on the ability to accurately detect individual cells over time based on image segmentation and are easily affected by such factors as plating density, image contrast variations, marker expression, low resolution image capture or occlusion, especially when the filming is over long time periods (e.g. a week). Particle image velocimetry (PIV) can extract motion in dense monolayers (25, 26) but subsequent downstream analysis do not systematically account for variable video quality, collective motion and global phenomena that occur across large groups of cells such as boundary formation. Previous analyses have only been applied to small video datasets and certain cell lines such as MDCK to measure specific motion properties such as speed which may not be discriminative for different motion phenotypes in general.

To address all the above informatics challenges we have developed Motion Sensing Superpixels (MOSES), a systematic motion analysis framework that does not require segmentation of individual cells. MOSES provides robust quantification of tissue-level dynamics. In contrast to existing methods it does not rely on lengthy data specific optimization of multiple parameters, specific image acquisition requirements or complex model fitting. Our computational framework provides a systematic approach to motion extraction, characterisation and phenotyping. We tested the potential of MOSES using a multi-well plate based *in-vitro* system at medium throughput to study the cell population dynamics and interactions between different epithelial cell types from the esophageal SCJ. The videos acquired were deliberately heterogeneous, acquired over separate days at varying imaging resolutions, different time lengths, assorted dye concentrations and various plating geometrics to test the ability of MOSES to handle challenging datasets. Our results demonstrate that MOSES robustly captures the relevant aspects of epithelial interactions and fulfils requirements for unbiased high content, comparative biological video analysis and phenotypic discovery.

## Results

### *In-vitro* model to study the spatio-temporal dynamics of boundary formation between different cell populations

To develop a motion analysis framework we established an experimental model to assess boundary formation between squamous and columnar epithelia found at the esophageal SCJ (Fig.1A). We used three epithelial cell lines: EPC2 (an immortalised squamous epithelial cell line from the normal esophagus (27)); CP-A (an immortalised BE cell line with columnar epithelial properties (28)); and OE33 (derived from EAC (29)). To model the interfaces that occur in the esophagus we used the combinations: EPC2:EPC2 (squamous:squamous, as a normal control), EPC2:CP-A (squamous:columnar, as in Barrett’s esophagus) and EPC2:OE33 (squamous:cancer, as in EAC) (Fig. 1B). In this experimental model (Fig.1C), two epithelial cell populations are co-cultured in the same well of a 24 well plate, separated by a divider with width 500µm. The divider is then removed after 12hr and cells allowed to migrate towards each other. Each cell population is labelled with a lipophilic membrane dye, which provides uniform staining, low phototoxicity and can be used *in-vivo* and to label primary cells (30). We tested the effects of the dye in two populations of EPC2, filmed over 96 and 144 hrs (Supp. Movie 1). The green and red fluorescent labelled EPC2 cells proliferated similarly, exhibiting the same cell density over the time course (slope=0.976, Pearson correlation coefficient 0.978; automatically counted from DAPI staining) (Fig.1D, Supp.Fig.1) and migrated similarly (assessed by the mean squared displacement (31)) (Supp. Fig 2). Thus, the dye has a minimal impact on cell motion behaviour. The effects of the dye were similarly checked in CP-A and OE33, (Fig. 1D, Supp.Fig.1D, 2 and Supp. Movie 1).

**Fig.1.**
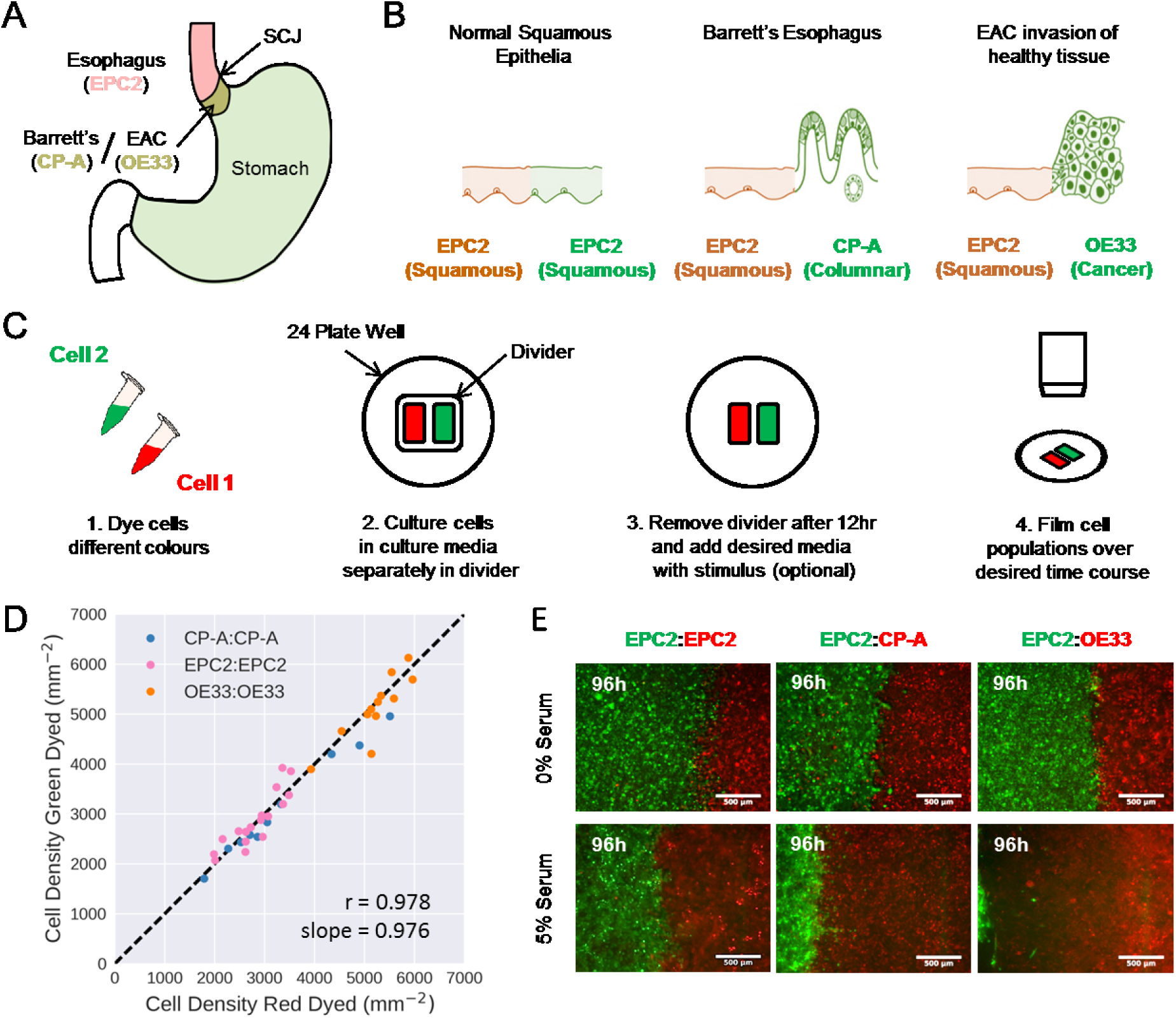
Temporary divider system to study interactions between cell populations. (*A*) The squamous columnar junction (SCJ) divides the stratified squamous epithelia of the esophagus and the columnar epithelia of the stomach. Barrett’s esophagus is characterised by squamous epithelia being replaced by columnar cells. (*B*) The three main epithelial interfaces that occur in BE to EAC progression. Pictures were adapted from figure 2 of (49). (*C*) Overview of the experimental procedure, described in steps 1-3. In our set-up cells were allowed to migrate and were filmed for 4-6 days after removal of the divider (step 4). (*D*) Cell density of red vs green dyed cells in the same culture, automatically counted from confocal images taken of fixed samples at 0,1,2,3,4 days and co-plotted on the same axis, (Supp. Fig. 1A,B for validation of the automatic counts with manual counts, Supp. Fig. 1D for counts plotted for each combination by day). Each point is a separate image. If a point lies on the identity line (black dashed), within the image, red and green dyed cells have the same cell density. (E) Snapshot at 96hr of three combinations of epithelial cell types, after culture without serum (upper) and with serum (lower). Scale bars: 500μm.

**Fig.2.**
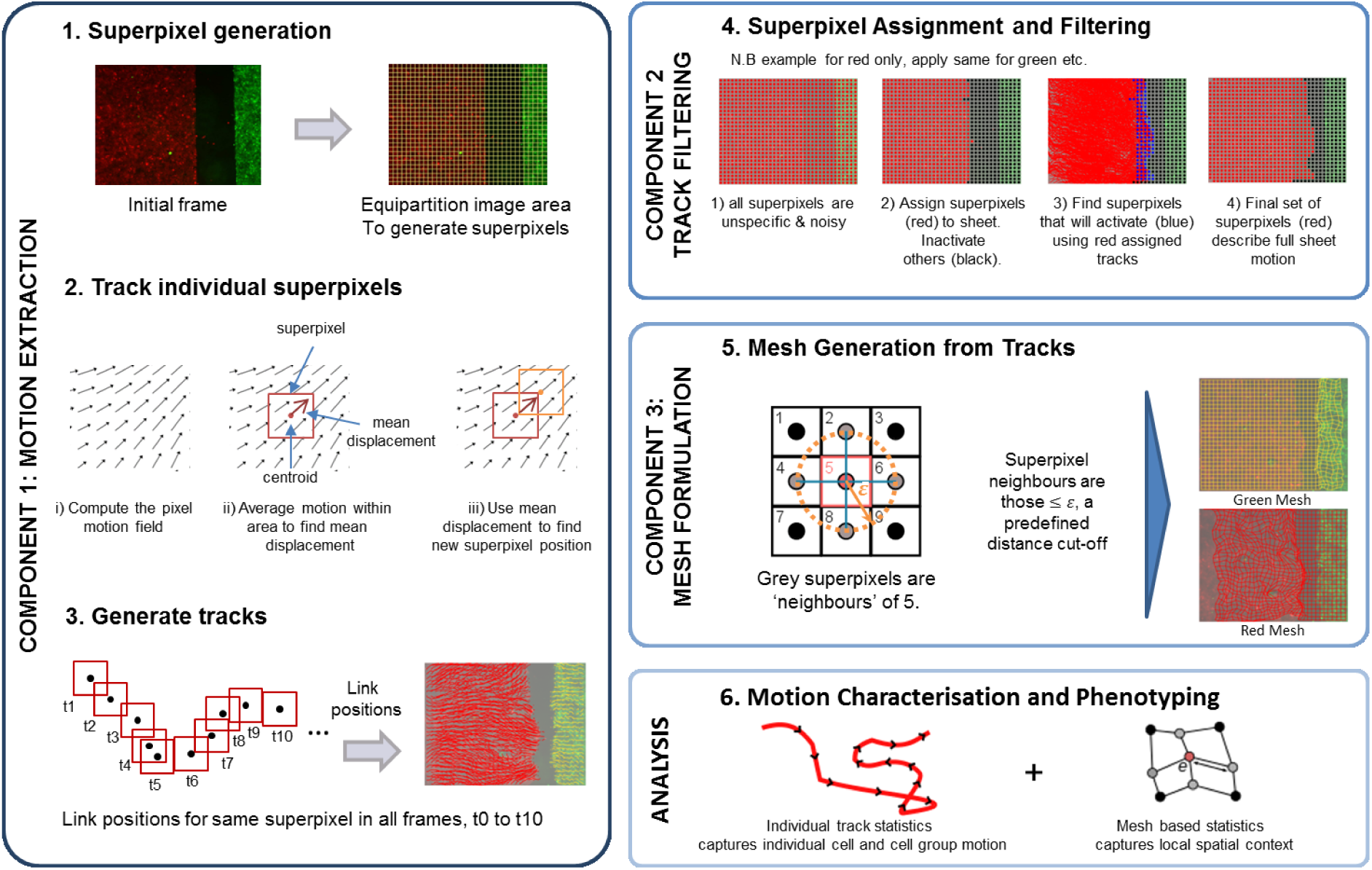
Schematic of MOtion SEnsing Superpixels (MOSES) framework for motion extraction and analysis. Motion extraction from a video is performed in steps 1-3; track filtering in step 4; and mesh formulation in step 5. Motion characterisation and phenotyping uses combined mesh and individual track statistics (step 6). See materials and methods and supporting information for detail.

We tested different combinations of the three epithelial cell lines, initially in serum-containing media (5% FBS) (Fig. 1E). In all combinations both populations moved as epithelial sheets (Supp. Movie 2). In the squamous EPC2:EPC2 combination, cells met and coalesced into a monolayer. In the squamous-columnar EPC2:CP-A combination a stable boundary formed between the two populations after 72 hrs, following a short period of CP-A pushing EPC2 (Supp. Movie 2, Fig. 1E). In the squamous:cancer EPC2:OE33 combination, the cancer cell line OE33 pushed EPC2 out of the field of view.

Evidence from systems including *Drosophila* embryonic parasegment (32) and anteroposterior and dorsoventral wing disc boundaries (33–35) suggest the importance of physical properties of cell/cell interactions for boundary formation and that collective migration is required for stable boundaries between epithelial populations. To test the importance of physical contact between cells in boundary formation, we cultured the same cell combinations in serum free medium to reduce cell-cell contacts (Supp. Fig. 3A). Under these conditions we observed loss of collective sheet migration (Supp. Fig. 3B,C) with no detectable boundary formation, and all cell combinations appeared to exhibit similar motion dynamics (Supp. Movie 3). This shows that cell-cell contacts are required for boundary formation. In subsequent experiments we used serum free conditions as a negative control for boundary formation.

**Fig.3.**
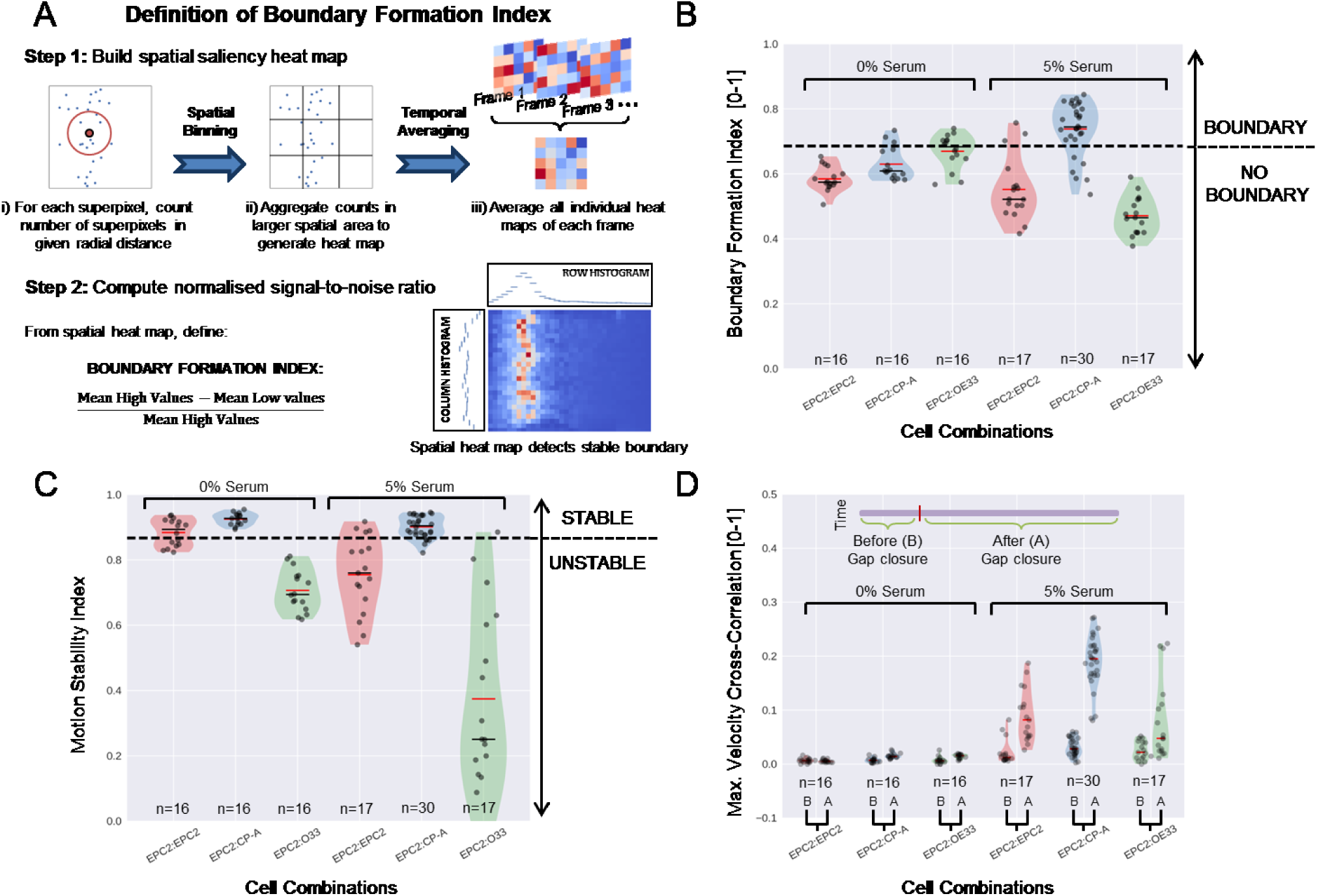
Quantitative assessment of boundary formation and sheet/sheet interaction dynamics. (*A*) Overview of the steps in calculating the boundary formation index. (*B*) Violin plots of boundary formation index for each video (black dot) for different cell combinations in 0% and 5% serum. Dashed line is the threshold given by one standard deviation above the pooled mean value of all cell combinations in 0% serum. Red solid line = mean, black solid line = median. Shaded region is the probability density of the data whose width is proportional to the number of videos at this value. (*C*) Motion stability index and (*D*) Maximum (Max.) velocity cross-correlation between the two sheets, shown as in (B) and as defined in SI materials and Methods. In (D), ‘B’ is before and ‘A’ is after gap closure.

### Development of MOSES to quantify cell motion dynamics

To address the computational challenges of analysing collective motion and cell interactions in a robust and sensitive manner we developed MOSES. The MOSES framework is modular and comprises three components (Fig.2): 1) motion extraction; 2) track filtering; and 3) “mesh” formulation. The first component is motion extraction from a video (steps 1-3, Fig.2) using superpixel tracking. As we cannot rely on the tracking of individual cells in tissue, this is done in a similar manner as the particle image velocimetry (PIV) approach for analysing monolayers and migrating epithelial fronts (e.g. (9)). Avoiding shortcomings of cell segmentation methods, the initial frame is divided into a specified number of regular regions to generate superpixels; each superpixel is larger than one image pixel (36) (Step1, Fig.2). The position of each superpixel is tracked frame-to-frame by updating its current (*x*,*y*) position according to the mean displacement calculated from optical flow (37) (Step 2, Fig.2). A superpixel track is generated by collecting the positions of a superpixel at all time points (Step 3, Fig.2). The collection of all superpixel tracks summarises all the motion within the video. The number of superpixels used should be chosen to sufficiently subsample the spatial area of the image to capture all spatial variations in motion. In this paper, we used 1000 superpixels to cover an image size of 1344 × 1024 (with this setting an average superpixel is 37 pixels × 37 pixels covering ≈ 14000μm^2^ (2x lens) or ≈57000μm^2^ (4x lens)) to track the sheet dynamics. The video channel for each different coloured cell population (i.e. red and green in our experimental system) is independently tracked in this manner.

As the superpixel tracks capture all the spatial motion dynamics in the video, some tracks may not be relevant to the phenomenon of interest. Therefore, the second component is track filtering (Step 4, Fig.2) and is specific to the experimental setup. In this step, superpixels are assigned to cover the entire dynamic motion for each ‘object’ of interest. For analysing epithelial sheets, the ‘object’ is the entire sheet; for single cell tracking the ‘object’ is each individual cell in the frame. To assign superpixels to an object, normally each object is first segmented from the initial video frame to produce a binary image where pixels have value 1 if belonging to the object or 0 otherwise. A superpixel then belongs to an object if its centroid lies in a pixel whose value is 1 in the respective binary image. However this approach is problematic for variable image intensities. To overcome this, we developed an intensity-independent segmentation method using the spatial layout of the superpixels for migrating epithelial sheets (Supp. Fig. 4, SI Materials and Methods).

**Fig.4.**
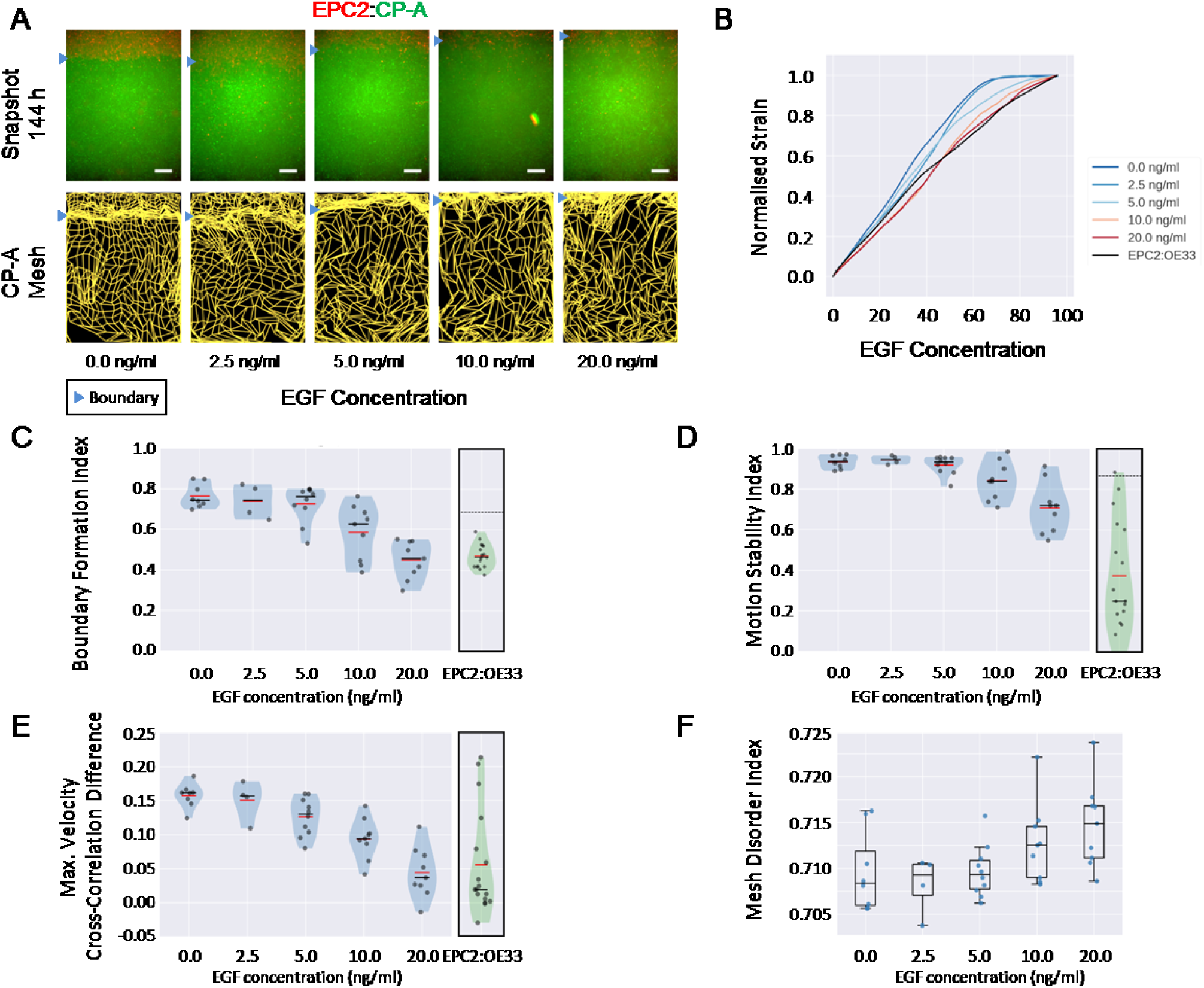
EGF titration at physiological levels disrupts boundary formation. (*A*) Destabilisation of the junction with EGF addition. All in 5% serum, snapshots at 144hrs of the green CP-A MOSES mesh at the same timepoint for all concentrations. The closeness of the lines indicates impeded motion leading to a local aggregation of superpixels in the vicinity and suggestive of a boundary. The less rectangular the mesh, the less ordered the motion. Blue triangles mark the boundary position in the image and its corresponding inferred position in the CP-A mesh. All scale bars: 500μm. (*B*) Mean normalised strain curves for each concentration. The mean curve for EPC2:OE33 videos in 5% serum without EGF in Fig. 3 is shown for comparison. Violin plots of boundary formation index (C), motion stability index (D) and maximum velocity cross-correlation (E) for each concentration of EGF. Red solid line = mean, Black solid line = median. Dots are individual videos, n=40. Values for EPC2:OE33 in 5% serum without EGF and threshold from Fig. 3 are shown for comparison. Shaded region is the probability density of the data whose width is proportional to the number of videos at this value. (*F*) Boxplot of mesh disorder index for each concentration of EGF.

The third component of MOSES is a “mesh formulation” to capture the spatial context of motion in tissue (Step 5, Fig.2). After filtering, spatial relationships between neighbouring superpixel tracks are captured in the form of a mesh. This transforms the independent trajectories of each superpixel into a dynamic mesh that naturally assimilates the local collective dynamics. A separate mesh is produced for each colour channel. Multiple different meshes can be constructed for different purposes and complex optimization is not required (see SI Materials and Methods). This mesh formalism is a key property of MOSES. Because the mesh is a representation of motion that is built from the local spatial relationships of superpixels, robust measures of motion relating to biological phenomena such as motion stability, collective motion and tissue interaction can be derived (see below). Analysis of motion characteristics and phenotype can use a combination of mesh-based statistics that account for the potential local collective motion, as well as single track statistics to measure the individual motion of cells (Step 6, Fig.2).

We tested the accuracy of the superpixel motion extraction experimentally by adding a sparse population of blue cells to the *in-vitro* setup described above (the blue cells being the same cell type as the population to which they were added) (Supp. Fig. 5A). We tested all three combinations of cell types (EPC2:EPC2, EPC2:CP-A and EPC2:OE33) and switched the red/green dye and the spiked-in population, generating 23 videos (each 144hr). To assess the performance of MOSES in tracking at particle level resolution (i.e. tracking the spiked-in cells), we used tracks generated by TrackMate (38), a published single particle tracker implemented in Fiji, as a benchmark. When the field of view was divided into 10000 superpixels in MOSES, the tracks were highly similar to TrackMate (Supp. Fig. 5B,C). To assess if the motion of single cells could be inferred from sheet motion, 1000 superpixels were used to track the motion of the sheet and the nearest superpixel tracks to each spiked-in cell were averaged to estimate the single cell trajectories and compared to TrackMate. Although the precise individual cell trajectory was lost, the overall motion pattern of the spiked in cells could be inferred (supp. Fig. 5B, C). Therefore, in confluent epithelial sheets where cells are ‘crowded’, individual cells behave similarly to their neighbours so global motion patterns can be used as a proxy to study single cell behaviour.

**Fig.5.**
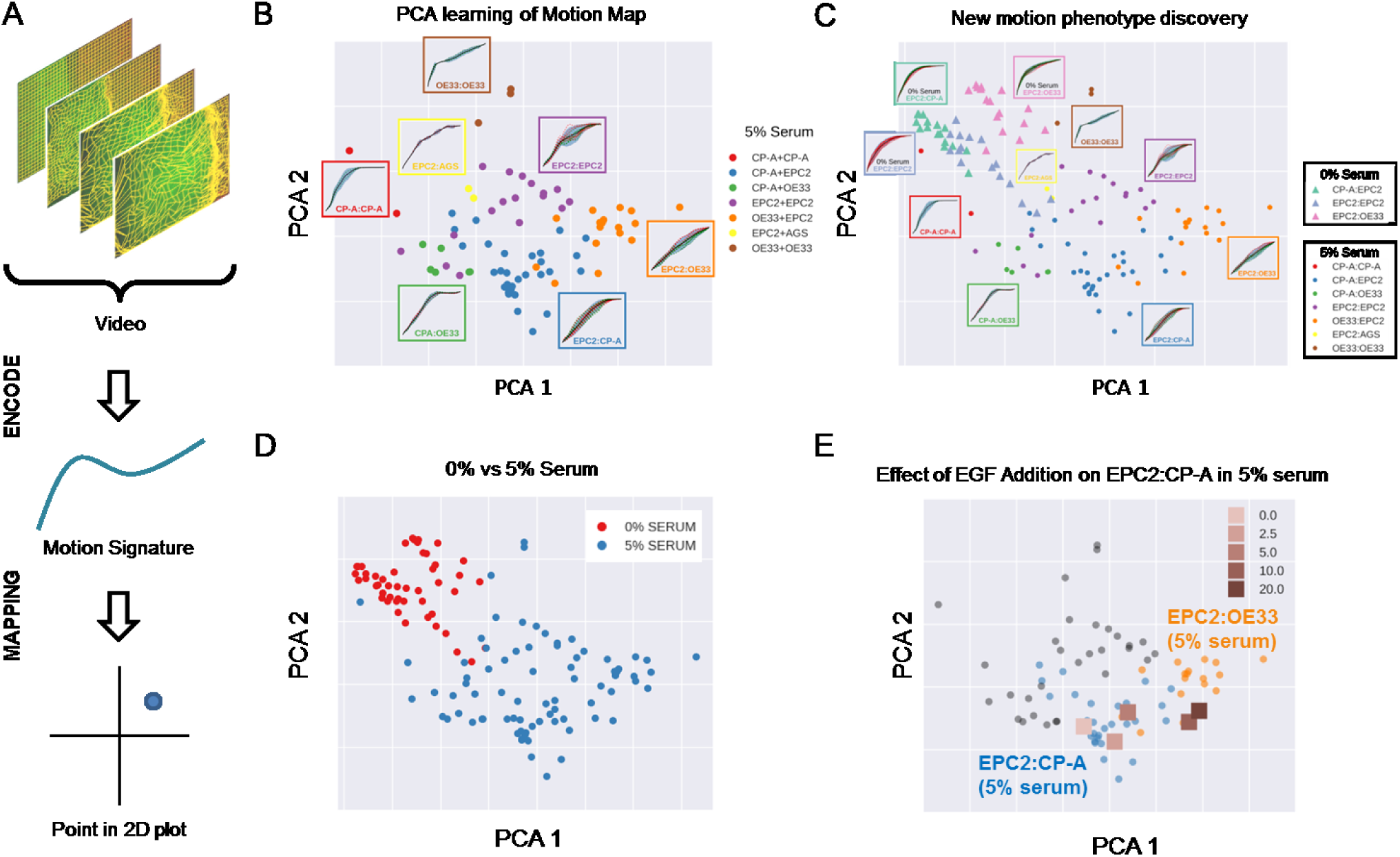
MOSES generates motion signatures to produce a 2D motion map for unbiased characterisation of cellular motion phenotypes. In all panels, each point represents a video (see legends for colour code). The position of each video on the 2D plot is based on the normalised mesh strain curves, analysed by PCA. (*A*) The mapping process for a single video. (*B*) The 5% serum videos (n=77) were used to set the PCA that maps a strain curve to a point in the 2D motion map. (*C*) The 0% serum videos (n=48) was plotted onto the same map defined by the 5% serum videos using the learnt PCA. In (*B*) and (*C*) the mean mesh strain curves for each cell combination are shown in insets. Light blue marks the two standard deviations with respect to the mean curve (solid black line). (*D*) Same map as in (*C*) with points coloured according to 0% or 5% serum. (*E*) The normalised mean strain curves for 0-20ng/ml EGF addition to EPC2:CP-A from Fig. 4B plotted onto the same map defined by the 5% serum videos.

### Quantitative measurement of squamous and columnar epithelial boundary formation using MOSES

To test the ability of MOSES to generate quantitative measurements that can be used to assess a specific biological feature, we used the example of stable boundary formation to define three indices: i) boundary formation index; ii) motion stability index; and iii) maximum velocity cross-correlation. Boundary formation and motion stability are mesh-derived statistics. Maximum velocity cross-correlation is derived from the individual superpixel tracks.

The boundary formation index is a global measure of the likelihood of stable boundary formation, by exploiting the notion that a boundary can be viewed as a spatially constrained phenomenon. By historically aggregating the number of neighbouring superpixels within a specified radius for each superpixel, we can generate a heatmap that spatially detects the interface from which we can quantify the global likelihood of boundary formation in a simple normalised signal to noise ratio index from 0-1, (Fig.3A); the higher the score the greater the likelihood of boundary formation.

The motion stability index is a local measure that assesses the additional stability of the interface between the two cell populations not captured by the boundary formation index, where individual cells in the sheet may still be moving stochastically without affecting the equilibrium spatial position and shape of the boundary. It is computed by subtracting from 1 the gradient of the normalised mesh strain curve, which measures the relative degree of motion of superpixels with respect to its neighbours within the tissue. The normalised mesh strain curve is a natural generalisation of the root mean squared displacement curve (RMSD) (materials and methods) to incorporate collective motion. The higher the motion stability index (maximum value of 1), the more stable the boundary and the cells at the boundary. The motion stability index for a cell combination is computed from the averaged normal mesh strain curve of the individual cell populations.

The maximum velocity cross correlation measures the degree that two epithelial sheets move together as a single cohesive connected sheet by correlating each superpixel track in one population (colour) with every superpixel track of the other population (colour), factoring in time delays. It is an averaged normalised measure (0-1). For two initially diametrically migrating sheets, a significant increase in this measure before and after closure of the gap is indicative of increased interaction across the sheets. This might be due to increased cell-cell contact in the combined monolayer leading to increased coordinated motion across larger spatial scales. These descriptors/measures are just examples of statistics that can be derived using MOSES. For further discussion and calculation of these metrics see SI materials and methods.

We used MOSES to compute these indices for different combinations of the three cell lines (EPC2, CP-A and OE33) in our experimental system. In total 125 videos (48 with 0% serum and 77 with 5% serum) were collected from 4 independent experiments and jointly analysed. These videos are highly heterogeneous and acquired under different conditions creating a challenging dataset for analysis (Supp. Table. 1, Supp. Fig. 6). Cells grown in 0% serum were used as negative control to set a cut-off for boundary formation (0.69) defined statistically as one standard deviation higher than the pooled mean of all three combinations (Fig.3B). Above this cut-off, cell combinations are categorised as forming a boundary. The boundary formation index was highest (0.74) for EPC2:CP-A grown in 5% serum (n=30/77) (Fig.3B, Supp. Fig. 7A). For EPC2:EPC2 and EPC2:OE33 in 5% serum the boundary formation index was below the cut-off (Fig.3B). We also ranked the serum videos on a continuous scale using the boundary formation index, which shows unbiasedly that the majority of EPC2:CP-A videos are at the top of this ranking (Supp. Fig. 8). Similarly, experiments in 0% serum were used to set the global motion stability threshold (0.87), one standard deviation below the pooled mean of EPC2:EPC2 and EPC2:CP-A. Cells at the interface of EPC2:OE33 are not stable and therefore not included in the pooled statistics. Below this cut-off, cell combinations are categorised as forming unstable interfaces. In 5% serum, EPC2:CP-A had the highest motion stability index with a median value of 0.90, compared to EPC2:EPC2 (0.76) and EPC2:OE33 (0.25) (Fig.3C, supp. Fig. 7B). These results illustrate that the squamous-columnar combination EPC2:CP-A forms a stable boundary.

We measured sheet-sheet interactions using the maximum velocity cross-correlation before and after gap closure for the three cell type combinations. In 0% serum there is no difference in velocity cross-correlation across all combinations before and after gap closure (Fig.3D). The two sheets do not move cohesively as a unit, which is to be expected with minimal cell-cell contact. In serum, there is greater cohesion due to cell-cell contact. We observe that the difference for EPC2:CP-A (median value of 0.03 before and 0.20 after gap closure) was ~3-6 times larger than for EPC2:EPC2 (0.01 to 0.08) and EPC2:OE33 (0.02 to 0.05), (c.f. left and right violins in Fig.3D). We note also that CP-A:OE33 (Barrett’s:cancer, n=6) also exhibited a substantial increase in velocity cross-correlation following gap closure (0.03 to 0.17) (Supp. Fig.7C,D). This is unlikely to be a feature of the CP-A cell line, as no substantial increase was observed for CP-A:CP-A (0.01 to 0.06) (Supp. Fig.7C,D). Thus, the EPC2:CP-A boundary exhibits greater cohesion between the two cell populations compared to interfaces formed between cells of the same type and EPC2:OE33 (i.e. ‘normal’ squamous cells with cancer cells) suggesting enhanced physical interaction. Of all the cell combinations tested, the squamous-columnar combination EPC2:CP-A uniquely forms an ‘interacting’ stable boundary.

### MOSES can measure subtle phenotype changes induced by external stimuli

For MOSES to be applicable for high-content imaging analyses, it needs to be able to quantitatively monitor subtle changes in cellular motion dynamics with a minimal number of replicates. As a test of the sensitivity of MOSES, we assessed whether it could detect subtle changes in the EPC2:CP-A boundary caused by an external stimulus.

The main cause of BE is bile acid reflux (39, 40). Bile acid activates epidermal growth factor receptor (EGFR) (41, 42), a receptor tyrosine kinase that is frequently mutated in EAC (43) and sometimes overexpressed in BE (44, 45). We therefore used EGF, the ligand of EGFR, as a stimulus to activate the EGFR signalling pathway. Increasing amounts of EGF (0ng/ml to 20ng/ml) were added to the culture medium to assess incremental effects on cellular motion and boundary formation in the EPC2:CP-A combination. A total of 40 videos (each 144hr) were collected from three independent experiments, in a 24-well plate medium throughput screen. Viewing the mesh (Fig. 4A), shows the boundary position further away from the initial point of contact between the two cell populations and decreased coherence of the boundary with increasing EGF. This is also shown by the shape of the mean normalised strain curve (Fig. 4B): at 0ng/ml EGF this curve linearly increases before plateauing around 72hrs; as EGF concentration increases, the curve becomes more linear and the plateau is lost above 5ng/ml. The boundary formation index decreases with increasing EGF (0.74 at 0ng/ml to 0.46 at 20ng/ml), indicating that the boundary is lost (i.e. index below the 0.69 cut-off) (Fig. 4C). The index at 20ng/ml EGF is similar to that for EPC2:OE33 without EGF (0.46), (Fig.4C). The interface becomes increasingly unstable and cells move more as the motion stability index decreases from 0ng/ml (0.94, stable) to 20ng/ml (0.72, unstable) (Fig.4D). Also, the interaction between the two cell populations is lost as the maximum velocity cross-correlation difference before and after gap closure decreased from 0ng/ml (0.16) to 20ng/ml (0.04) EGF (Fig. 4E). The maximum velocity cross-correlation after gap closure is similar to that for EPC2:OE33 (0.02) (Fig.4C), but the motion stability index remains higher (Fig.4D). Altogether these measures show that above 5 ng/ml EGF the phenotype of EPC2:CP-A becomes similar to that of the interaction between EPC2 and the EAC cell line OE33 (supp. Movie 4).

We also developed an index of the level of disorder in the MOSES mesh (SI materials and methods) (Supp. Fig. 9A), which captures the extent local cell populations move in opposing directions with respect to their neighbours. The mesh disorder index showed statistically significant increases with EGF concentration (0ng/ml: 0.708, 2.5ng/ml: 0.709, 5ng/ml: 0.709, 10ng/ml: 0.713, 20ng/ml: 0.715) (Fig.4F, Supp. Fig. 9B,C), suggesting collective sheet motion is lost. This loss of collective motion is supported by a decrease in the standard spatial correlation measure (SI materials and methods). However, using this standard approach gives high statistical variance, (Supp. Fig. 9D). The effect of increased disorder is visually subtle (supp. Movie. 4), but is clearly detected using the MOSES mesh and the proposed mesh disorder index.

Titrating EGF in the absence of serum gave non-significant changes in the boundary formation index (0ng/ml: 0.60±0.07, 20ng/ml: 0.65±0.03) and maximum velocity cross-correlation (supp. Fig.10 (n=25)). Decreasing motion stability index indicates increased cell movement and increasing mesh disorder index suggests the absence of collective dynamics required for boundary formation. In summary, this example with EGF in the context of our experimental set-up shows that MOSES enables continuous-scale quantification of motion after systematic perturbation in a medium-throughput 24-well format.

### MOSES generates motion signatures and 2D motion maps for unbiased characterisation of cellular motion phenotypes

We have shown how the mesh-based formulation of MOSES can be used to derive robust measurements to test directly for differences in expected motion phenotypes, such as boundary formation. But what if we do not know what motion phenotype to expect? For example, for high content imaging screens it is necessary to be able to screen for unknown differences in complex cellular motions from a large numbers of videos in an unbiased manner. MOSES addresses this need by facilitating the systematic generation of unique ‘motion signatures’ for each video. Here we demonstrate that unsupervised machine learning techniques requiring no manual user annotation can be used to plot videos onto a 2D motion phenotype map, enabling easy visual assessment of motion phenotype and hypothesis generation without the need to individually interrogate each video.

The general process for motion map generation is illustrated in Fig. 5A. For each video to be positioned on a 2D map, we applied principal components analysis (PCA) to the normalised mesh strain curves for the 77 videos of cell combinations cultured in 5% serum conditions. The mesh strain curve for each video (used above to define the motion stability index) is used here as a 1D motion signature for summarising the entire video motion, (see SI material and methods for generating other descriptive feature vectors). The map generated for these 77 videos (Fig. 5B) shows that this unbiased approach has automatically clustered together the videos for each cell type combination. Furthermore, the videos are ordered in a continuous manner, as shown by the progressive transformation in the shape of the mean normalised strain curve when looking across the plot in Fig. 5B from left to right, CP-A:CP-A to EPC2:OE33 (i.e. the shape is increasingly linear). This result could not be achieved with RMSD (Supp. Fig. 11), and is independent of the particular dimensionality reduction technique used, (Supp. Fig. 12). Further, the 1D motion signatures derived from MOSES could be used to train a machine learning classifier with no further processing to predict cell combination identity better than RMSD, (Supp. Fig. 11).

To demonstrate how 2D motion phenotype maps generated by MOSES can be used to compare videos, we next mapped the 48 videos from 0% serum cultures on the same axes as the videos from 5% serum (Fig. 5C,D). The videos from 0% serum mapped to a different area of the 2D plot, whilst preserving the continuous ordering of the previous videos. Therefore, without having watched the videos it is easy to predict that the cells have markedly different motion dynamics in 0% serum compared to 5% serum. Furthermore, since the points for the 5% serum videos cover a larger area of the 2D plot than the 0% serum videos, one would expect less diversity of motion in 0% serum, (Fig.5D).

The motion map can also capture subtle motion changes. This is demonstrated by mapping the mean video motion for each concentration of EGF from 0-20ng/ml (represented by the respective mean normalised strain curves for each concentration (1 per concentration from a total n=40 videos, see Fig.4B)) onto the same axis as the 5% serum videos in the absence of EGF (square points in Fig 5E). With increasing EGF, the EPC2:CP-A motion dynamics become more similar to EPC2:OE33, as evidenced above 5ng/ml by the square points moving from the area of blue circular EPC2:CP-A points into the area of orange circular EPC2:OE33 points. Therefore, the motion map is consistent with the results obtained using the specific derived indices (above). These results illustrate that MOSES is able to account for biological and technical variability across independent experiments and possesses the required features for an algorithm to be used in an unbiased manner in a high content screening.

## Discussion

MOSES satisfies the four criteria necessary for characterising and establishing phenotypes from live cell imaging. By analysing highly heterogeneous videos the mesh formulation of MOSES demonstrates potential to analyse challenging datasets in situations (such as *intra-vital* imaging or high throughput screening) where image quality cannot be ideally controlled, number of replicates is low, poor quality videos cannot be easily discarded, and for which alternative methods may be too time-consuming. Different components of MOSES can also be adapted to suit different applications, such as non-square superpixel shapes to better capture cells that undergo large shape changes.

The accuracy of the superpixel motion extraction was compared to a published single-cell tracker and demonstrated robust recovery of single cell-level tracks from global motion. This experimentally and quantitatively justifies the widespread use of global tracking such as PIV analysis for studying cellular motion in confluent monolayers where accurate individual cell segmentation is not possible. Previously the use of PIV has only been justified indirectly by comparing derived measurements, such as speed (25), or by analogy of the epithelial sheet motion to fluid-like dynamics (26, 46), or reasoned from force measurements (47) or qualitative comparison (48).

MOSES does not require complex user settings to facilitate reproducibility in analyses. The main parameter the user needs to specify is the number of initial superpixels. No complicated fitting of complex models and no special hardware is required. It can be run on a single PC (3.2Ghz, 16Gb RAM). Analysis of 96 videos with pixel resolution 1344×1024, 145 frames, tracking 1000 superpixels is under 4hr with results stored efficiently (~1-2 Mb per video). MOSES could be useful for screening the effects of drugs in a setting with ‘target’ cells (e.g. cancer cells) and other cells (e.g. representing adjacent normal tissue), to provide a more physiologically relevant system for screening effects on specific cell types and their interactions. Finally MOSES can be readily incorporated into existing motion analytical workflows based on single cell and PIV tracking.

## Materials and Methods

### Cell lines and Tissue Culture

EPC2 and CP-A (ATCC) cells were grown in full KSFM (Thermo Fisher), AGS (ATCC) and OE33 (ATCC) in full RPMI with 10% FBS. Both were supplemented with glutamine and Penicillin streptomycin at 37^0^C and 5% CO_2_ until 80% confluent. To passage EPC2 and CP-A, cells were resuspended after trypsinization for 5 min with PBS supplemented with soybean trypsin inhibitor (0.25 g/L, Sigma) to prevent cell death prior to resuspension in KSFM. To store, cells were resuspended at a concentration of 10^6^ cells/mL with 90% FBS + 10% DMSO freezing media following centrifugation and stored at −80°C before passing to liquid nitrogen storage.

### Fluorescent Labelling

Cells were labelled using Celltracker Green CMFDA and Celltracker Orange CMRA dyes (Life Technologies) according to protocol. Two different concentrations 2.5µM and 10µM were used. The lower concentration still permits tracking with less toxicity concerns.

### Temporary Divider Co-culture Assay

70,000 labelled cells in 70µL of culture media were seeded into each side of a cell culture insert (Ibidi). After 12h inserts were removed and the well washed three times with PBS to remove non-attached cells before adding the desired media (KSFM (0% serum in the text) or 1:1 mixture of KSFM:RPMI + 5%FBS (5% serum in the text)) for filming. For the perturbations the effector, e.g. EGF, was also added to the media at the stated concentrations.

### Image Acquisition

The different conditions were filmed on a Nikon microscope for 96 or 144 h at a frequency of 1 image per hour. 2x and 4x objectives were used. 546nm was used for the red dye and 488nm for the green dye.

### MOSES Framework

MOSES was developed using the Python Anaconda 2.7 distribution, in particular it uses Numpy, the Scipy-stack and OpenCV libraries. It comprises separately a cell tracking and data analysis component.

### Motion Extraction

Regular superpixels were generated by applying the SLIC (36) algorithm in scikit-image on a blank image, the same size as the video frame. 1000 superpixels were used throughout the paper. Motion fields for updating the superpixel centroid position over time were computed with Farneback optical flow (37) in OpenCV using default parameters. For ease of implementation displacement vectors were rounded to the nearest integer. Superpixels passing out of the frame progressively lose pixels and retain their last motion position for the tracking duration. For simplicity lost area is not recovered.

### Dynamic Mesh Generation

To generate meshes connect each superpixel to its nearest ‘neighbours’. For the time-dependent strain MOSES mesh used for visualisation and stability analysis, the ‘neighbours’ are determined only by which superpixels were closer than a specified distance at the start of tracking. For computation of the boundary formation index for each frame, neighbours were independently determined frame-by-frame by a preset distance threshold. The specified threshold in both cases is given in Euclidean distance as a multiplicative factor of the superpixel width for that video. A factor of 1.2 for the mesh strain and 5.0 was used for the junction formation index throughout.

### Normalised Mesh Strain Curves

For a superpixel at time *t* we define the local neighbourhood *N* mesh strain *ε(t)* with *n* neighbours as the mean of the distances *x_j_(t)* from the equilibrium position *x_j_*(0) at the start *t=0* so that 
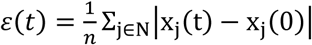
 where | ⋅ | is the *L1*-norm. We use *L1* for robustness as this value is > 1. The time-dependent mesh strain for one mesh is the mean local neighbourhood mesh strain over all superpixels for each time frame. The result is a vector the same length as the number of frames in the video. For multi-channels the average vector is used to describe video motion. The absolute value of the strain curve is susceptible to the image acquisition conditions and geometry whilst the motion information is primarily encoded by the shape of the resulting curve. To permit comparison across different conditions the normalised strain curve (division by maximum strain) was used as a simple signature to describe the global video motion pattern.

### Mean Squared Displacement (MSD)

As a measure of cellular motions, MSD was computed as a function of time interval, Δ*t* as in (31). MSD 
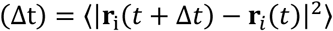
 where r*_i_(t)* is the position of the superpixel *i* at time *t* and 〈 ⋅ 〉 is the average over all time *t* and all superpixels. For small Δ*t*, the MSD increases as a power law Δ*t^α^*, where the exponent α is determined empirically by fitting. For unity exponent (*α* = 1) the movement is uncorrelated random Brownian motion and cellular motion is diffusive. When *α* > 1, cellular motions are super-diffusive, and when *α* = 2, motions are ‘ballistic’.

### Root Mean Squared Displacement (RMSD)

As a summary of the whole video motion, the root mean squared displacement was computed as a function of time relative to the initial time *t*_0_, RMSD 
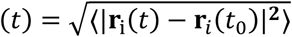
 where **r***_i_(t)* is the position of the superpixel *i* at time *t* and 〈 ⋅ 〉 is the average over all time *t* and all superpixels.

### Quantification Metrics

Details for the computation of the boundary formation, motion stability, normalised velocity correlation and mesh disorder measures are available in the supporting information materials and methods.

## Software Availability

Example data for testing the code can be downloaded from the following Google Drive folder with the hyperlink, https://drive.google.com/open?id=0BwFVL6r9ww5BaTh6NExLR1JMUXM. Software code is made available on GitHub (https://github.com/fyz11/MOSES.git).

## Conflict of Interest

A patent is pending for MOSES. The current code version is available open-source and free for academic and non-profit users under a Ludwig Software License. Details in the Github.

## Author Contributions

F.Y.Z conceived and implemented MOSES. C.R.P carried out all live cell imaging videos and established the 24-well plate based *in-vitro* co-culture systems. R.P.O, M.J.W aided initial prototyping of MOSES with mixed co-culture timelapse videos. J.R provided computational supervision and X.L provided overall supervision of the project. The manuscript was predominantly written by F.Y.Z and X.L, with input from all authors.

## Acknowledgements

We thank Prof Hiroshi Nakagawa for his generous donation of EPC2 cells. We thank Dr Mary Muers, Profs. Sebastian Nijman, Francis Szele, Colin Goding and Shankar Srinivas for critical reading and suggestions of the manuscript; Mark Shipman for technical assistance with timelapse microscopy. This work is mainly funded by the Ludwig Institute for Cancer Research (LICR) with additional support from a CRUK grant to XL. F.Y.Z is mainly funded through the EPSRC life sciences interface doctoral training centre EP/F500394/1, C.R.P and XL are funded by LICR, R.P.O is supported by the NIHR Biomedical Research Centre, Oxford, LICR, CRUK, OUCAG, M.J.W. is supported by CRUK, and JR is funded by LICR and the EPSRC SeeBiByte Programme Grant (EP/M013774/1).

## Bibliography

1. Schaffer CJ & Nanney LB (1996) Cell biology of wound healing. International review of cytology 169:151–181.

2. Clark R (2013) The molecular and cellular biology of wound repair (Springer Science & Business Media).

3. Leoni G, Neumann P, Sumagin R, Denning T, & Nusrat A (2015) Wound repair: role of immune–epithelial interactions. Mucosal immunology 8(5):959–968.

4. Dahmann C, Oates AC, & Brand M (2011) Boundary formation and maintenance in tissue development. Nature Reviews Genetics 12(1):43–55.

5. Gaddam S, et al. (2013) Persistence of nondysplastic Barrett’s esophagus identifies patients at lower risk for esophageal adenocarcinoma: results from a large multicenter cohort. Gastroenterology 145(3):548–553. e541.

6. Schmitz MH, et al. (2010) Live-cell imaging RNAi screen identifies PP2A-B55 [alpha] and importin-[beta] 1 as key mitotic exit regulators in human cells. Nature cell biology 12(9):886–893.

7. Pau G, et al. (2013) Dynamical modelling of phenotypes in a genome-wide RNAi live-cell imaging assay. BMC bioinformatics 14(1):308.

8. Held M, et al. (2010) CellCognition: time-resolved phenotype annotation in high-throughput live cell imaging. Nature methods 7(9):747–754.

9. Das T, et al. (2015) A molecular mechanotransduction pathway regulates collective migration of epithelial cells. Nature cell biology 17(3):276–287.

10. Togashi H, et al. (2011) Nectins establish a checkerboard-like cellular pattern in the auditory epithelium. Science 333(6046):1144–1147.

11. Javaherian S, et al. (2015) An in vitro model of tissue boundary formation for dissecting the contribution of different boundary forming mechanisms. Integrative Biology 7(3):298–312.

12. Fletcher AG, Osterfield M, Baker RE, & Shvartsman SY (2014) Vertex models of epithelial morphogenesis. Biophysical journal 106(11):2291–2304.

13. Alt S, Ganguly P, & Salbreux G (2017) Vertex models: from cell mechanics to tissue morphogenesis. Phil. Trans. R. Soc. B 372(1720):20150520.

14. Cai AQ, Landman KA, & Hughes BD (2007) Multi-scale modeling of a wound-healing cell migration assay. Journal of Theoretical Biology 245(3):576–594.

15. Markham DC, Simpson MJ, & Baker RE (2015) Choosing an appropriate modelling framework for analysing multispecies co-culture cell biology experiments. Bulletin of mathematical biology 77(4):713–734.

16. Podewitz N, Jülicher F, Gompper G, & Elgeti J (2016) Interface dynamics of competing tissues. New Journal of Physics 18(8):083020.

17. Hatzikirou H & Deutsch A (2008) Cellular automata as microscopic models of cell migration in heterogeneous environments. Current topics in developmental biology 81:401–434.

18. Mallet DG & De Pillis LG (2006) A cellular automata model of tumor–immune system interactions. Journal of Theoretical Biology 239(3):334–350.

19. Meijering E, Dzyubachyk O, & Smal I (2012) 9 Methods for Cell and Particle Tracking. Methods in enzymology 504(9):183–200.

20. Masuzzo P, Van Troys M, Ampe C, & Martens L (2016) Taking aim at moving targets in computational cell migration. Trends in cell biology 26(2):88–110.

21. Padfield D, Rittscher J, & Roysam B (2011) Coupled minimum-cost flow cell tracking for high-throughput quantitative analysis. Medical image analysis 15(4):650–668.

22. Maška M, et al. (2014) A benchmark for comparison of cell tracking algorithms. Bioinformatics 30(11):1609–1617.

23. Hilsenbeck O, et al. (2016) Software tools for single-cell tracking and quantification of cellular and molecular properties. Nature biotechnology 34(7):703–706.

24. Schiegg M, et al. (2015) Graphical model for joint segmentation and tracking of multiple dividing cells. Bioinformatics 31(6):948–956.

25. Petitjean L, et al. (2010) Velocity fields in a collectively migrating epithelium. Biophysical journal 98(9):1790–1800.

26. Szabo B, et al. (2006) Phase transition in the collective migration of tissue cells: experiment and model. Physical Review E 74(6):061908.

27. Harada H, et al. (2003) Telomerase Induces Immortalization of Human Esophageal Keratinocytes Without p16INK4a Inactivation1 1 NIH grants R01-DK5337 (AKR), P01-DE12467 (AKR), P01-CA098101 (AKR), Deutsche Krebshilfe 10-1656-Op 1 and D/96/17197 (OGO), NIH R21 DK64249-01 (HN), AGA/FDHN Fiterman Award and American Cancer Society (GHE), and NIH/NIDDK Center for Molecular Studies in Digestive and Liver Diseases (P30 DK50306). Molecular Cancer Research 1(10):729–738.

28. Merlo LM, Kosoff RE, Gardiner KL, & Maley CC (2011) An in vitro co-culture model of esophageal cells identifies ascorbic acid as a modulator of cell competition. BMC cancer 11(1):461.

29. Boonstra JJ, et al. (2010) Verification and unmasking of widely used human esophageal adenocarcinoma cell lines. Journal of the National Cancer Institute 102(4):271–274.

30. Progatzky F, Dallman MJ, & Celso CL (2013) From seeing to believing: labelling strategies for in vivo cell-tracking experiments. Interface focus 3(3):20130001.

31. Park J-A, et al. (2015) Unjamming and cell shape in the asthmatic airway epithelium. Nature materials 14(10):1040–1048.

32. Monier B, Pélissier-Monier A, Brand AH, & Sanson B (2010) An actomyosin-based barrier inhibits cell mixing at compartmental boundaries in Drosophila embryos. Nature cell biology 12(1):60–65.

33. Landsberg KP, et al. (2009) Increased cell bond tension governs cell sorting at the Drosophila anteroposterior compartment boundary. Current Biology 19(22):1950–1955.

34. Major RJ & Irvine KD (2005) Influence of Notch on dorsoventral compartmentalization and actin organization in the Drosophila wing. Development 132(17):3823–3833.

35. Major RJ & Irvine KD (2006) Localization and requirement for Myosin II at the dorsal‐ ventral compartment boundary of the Drosophila wing. Developmental dynamics 235(11):3051–3058.

36. Achanta R, et al. (2012) SLIC superpixels compared to state-of-the-art superpixel methods. IEEE transactions on pattern analysis and machine intelligence 34(11):2274–2282.

37. Farnebäck G (2003) Two-frame motion estimation based on polynomial expansion. Scandinavian conference on Image analysis, (Springer), pp 363–370.

38. Tinevez J-Y, et al. (2017) TrackMate: An open and extensible platform for single-particle tracking. Methods 115:80–90.

39. Souza RF (2010) The role of acid and bile reflux in oesophagitis and Barrett’s metaplasia. (Portland Press Limited).

40. Dixon M, Neville P, Mapstone N, Moayyedi P, & Axon A (2001) Bile reflux gastritis and Barrett’s oesophagus: further evidence of a role for duodenogastro-oesophageal reflux? Gut 49(3):359–363.

41. Werneburg NW, Yoon J-H, Higuchi H, & Gores GJ (2003) Bile acids activate EGF receptor via a TGF-α-dependent mechanism in human cholangiocyte cell lines. American Journal of Physiology-Gastrointestinal and Liver Physiology 285(1):G31–G36.

42. Dossa AY, et al. (2015) Bile acids regulate intestinal cell proliferation by modulating EGFR and FXR signaling. American Journal of Physiology-Gastrointestinal and Liver Physiology:ajpgi. 00065.02015.

43. Secrier M, et al. (2016) Mutational signatures in esophageal adenocarcinoma define etiologically distinct subgroups with therapeutic relevance. Nature Genetics 48(10):1131–1141.

44. Cronin J, et al. (2011) Epidermal growth factor receptor (EGFR) is overexpressed in high-grade dysplasia and adenocarcinoma of the esophagus and may represent a biomarker of histological progression in Barrett’s esophagus (BE). The American journal of gastroenterology 106(1):46–56.

45. Al‐Kasspooles M, Moore JH, Orringer MB, & Beer DG (1993) Amplification and over‐ expression of the EGFR and erbB‐2 genes in human esophageal adenocarcinomas. International journal of cancer 54(2):213–219.

46. Angelini TE, Hannezo E, Trepat X, Fredberg JJ, & Weitz DA (2010) Cell migration driven by cooperative substrate deformation patterns. Physical review letters 104(16):168104.

47. Trepat X, et al. (2009) Physical forces during collective cell migration. Nature physics 5(6):426–430.

48. Zaritsky A, Natan S, Ben-Jacob E, & Tsarfaty I (2012) Emergence of HGF/SF-induced coordinated cellular motility. PLoS One 7(9):e44671.

49. Evans RP, Mourad MM, Fisher SG, & Bramhall SR (2016) Evolving management of metaplasia and dysplasia in Barrett’s epithelium. World journal of gastroenterology 22(47):10316.

